# Contrasting drivers of abundant phage and prokaryotic communities in tropical, coastal ecosystems across the Isthmus of Panama

**DOI:** 10.1101/2023.01.26.525695

**Authors:** Alaina R. Weinheimer, Frank O. Aylward, Matthieu Leray, Jarrod J. Scott

## Abstract

Phages, or viruses that infect bacteria and archaea, are ubiquitous and abundant members of Earth’s ecosystems that impact the flow of nutrients, evolution of microbes, and food web dynamics by selectively infecting and killing their prokaryotic hosts. Because phages can only replicate through their hosts, they are inherently linked to processes impacting their hosts’ distribution and susceptibility to infection. Despite these links, phages can also be affected by environmental parameters independent of their hosts, such as pH or salinity which impact cell adsorption or virion degradation. To understand these complex links, in this study, we leverage the unique ecological context of the Isthmus of Panama, which narrowly disconnects the productive Tropical Eastern Pacific (TEP) and Tropical Western Atlantic (TWA) provinces and compare factors that shape active marine phage and prokaryotic communities. Metagenomic sequencing of seawater from mangroves and reefs of both the TEP and TWA coasts of Panama suggest that pronounced environmental gradients, such as along the TEP mangrove rivers, result in common dispersal and physicochemical parameters shaping both prokaryotic and phage community composition and diversity. Conversely, we find that when environmental conditions are relatively similar across adjacent habitats, such as between the mangroves and reefs in the TWA, prokaryotic communities are more influenced by local abiotic conditions while phage communities are shaped more by dispersal. Collectively, this work provides a framework for addressing the co-variability between viruses and their hosts in marine systems and for identifying the different factors that drive consistent versus disparate trends in community shifts, which is essential to inform models of these interactions in biogeochemical cycling.

## INTRODUCTION

Microbes are crucial components of Earth’s ecosystems, particularly in the ocean, where they form the foundation of food webs, power biogeochemical cycles, and expand the ecological niches of plants and animals^1,2^. Outnumbering even microbes, viruses serve as major top-down control on microbial communities and modulate microbial ecology and evolution through selective killing via infections, horizontal gene transfer via transduction, and metabolic reprogramming during infections^3^. Understanding viral impacts on microbes is critical toward modeling the movement of nutrients through ecosystems^4^, the evolution of microbial pathogens^5^, and the dynamics of organismal-associated microbiomes^6^. While rapid advances in sequencing and microscopy technologies over the past few decades have begun to unfold the vast diversity, complexity, and breadth of viruses in nature^7–9^, major questions remain on which factors shape viral communities and how this relates to concomitant shifts in microbial communities.

Because viruses are restricted to reproducing through their hosts, viruses are inherently linked to processes related to their hosts’ distribution and susceptibility to infection. Despite these tight links, patterns in the composition and diversity of these two groups can differ depending on the parameter or environment. Showing coupled shifts, for instance, viral diversity and microbial diversity in the ocean has been shown to increase with depth^10,11^, and the pH of soils has been shown to co-vary with viral and prokaryotic (bacteria or archaea) diversity^12^. In contrast to this coupling, a study examining soil communities and one on communities in freshwater springs showed that viral communities shifted over spatial scales and environmental parameters that did not always match that of microbes^13,14^. Several possibilities have been suggested to explain these contrasts, such as a broader host range of viruses lowering the impact of available host composition on viral community structure^15^, or metacommunity dynamics^14^ such as the importance of high dispersal versus species local adaptation that may differ between microbes and viruses. Taken together, these studies highlight the necessity to untangle the complexity in the link between viral communities and microbial communities, to better characterize roles of microbes and viruses in the environment.

In this study, we leverage the unique biogeography of the Isthmus of Panama to uncover factors shaping viral and microbial communities across a diverse array of tropical coastal environments in two oceans. The Isthmus of Panama gradually formed and finally completely disconnected the Tropical Western Atlantic Ocean (TWA) from the Tropical Eastern Pacific Ocean (TEP) approximately 2.8 million years ago^16^. The TWA became oligotrophic, leading to the proliferation of reef-building corals. The TEP remained eutrophic, with patchy coral reefs dominated by fewer species of scleractinian corals. Expansive mangroves thrive adjacent to coral reefs in both the TEP and the TWA. Nonetheless, mangroves of the TWA are influenced by much smaller tidal oscillations than in the TEP. In addition, the TWA supports thinner fringes of mangroves made of shorter trees than in the productive TEP^17^. These contrasting coasts with parallel habitat types of mangroves and coral reefs allow comparisons of viral and microbial communities at two spatial scales, locally between habitat types and globally between oceans^18^. Given the intrinsic link of viruses to their hosts, our null hypothesis was that factors shaping viral communities mirror those of microbial communities, and this similarity would be most visible at global scales between the oceans since the spatial separation and chemical differences between oceans are so large. An alternative hypothesis is that factors shaping viral communities would not match those of the corresponding microbial communities, and these differences would be most apparent at smaller scales where subtle differences in environmental parameters can influence contact rates of viruses to hosts, growth rates of hosts, and other physical aspects that may decouple viral communities from microbial communities.

To address these hypotheses, we examined both prokaryotic and viral community diversity in seawater metagenomes filtered for the 0.22-0.8 μm size fraction. While most known viruses are smaller than 0.22 μm, the viruses detected in this size fraction correspond to a subset of the viral community that includes larger viruses (e.g. jumbo bacteriophages), actively replicating (pre-lytic) viruses, lysogenic viruses (those integrated in the genomes of hosts) or viruses that have stuck to particles, putatively representing an active or abundant subset of the viral community^19^. Because viruses of bacteria and archaea that belong to the class *Caudoviricetes* are ubiquitous members of ecosystems, we focused our analyses on these viruses that we refer to as phages. For the microbes, we focused on the prokaryotes, as they are the putative host pool of the phages. To directly compare phage and prokaryotic diversity and minimize information loss, we applied a gene-based approach. We selected marker genes for both prokaryotes and phages that we benchmarked against more commonly used contig-based analyses.

Our results reveal a variety of contexts when factors shaping phage and prokaryotic communities align and when they diverge. Supporting our null hypothesis, the phage and prokaryotic communities were both distinct between oceans. The importance of habitat type, however, differed between these groups. Distinctions in phage community composition between mangroves and reefs depended on the ocean, with the mangrove communities being highly distinct from the reef communities in the TEP but not in the TWA, likely due to the strong salinity gradient of the mangrove rivers in the TEP. Meanwhile, the prokaryotic communities were equally distinct between the habitat types in both oceans. The lack of separation between the habitat types of the phage communities in the TWA compared to the prokaryotic communities suggests that changes in environmental parameters influence prokaryotic communities nearly equally as dispersal limits or physical separation, while phage communities are more structured by dispersal limitations than local conditions. Most strikingly, we found that phage communities were more diverse in the TWA, while prokaryotic communities were more diverse in the TEP, suggesting phage production or breadth of host range may differ in ecosystems of the more productive island archipelago of the TEP than in the oligotrophic coastal bay of the TWA. Overall, these findings highlight the necessity to examine viruses with their potential host community together to better untangle processes driving their interactions with each other and the environment in natural, mixed communities. The contexts when phage and prokaryotic communities do not couple each other is crucial for modeling phage-host interactions as they relate to microbial mortality, and ultimately biogeochemical cycling in ecosystems.

## RESULTS AND DISCUSSION

### Benchmarking methods to assess phage and prokaryotic diversity

In total, fifty-seven samples of seawater from mangroves and reefs were collected from the TWA and TEP coasts of Panama for metagenomic sequencing (Figure 1). To directly compare phage and prokaryotic diversity and minimize information loss from the metagenomic data, we benchmarked and employed a novel gene-based approach (see Methods), in which families of the major capsid protein (MCP) and terminase large subunit (TerL) belonging to the class *Caudoviricetes* compiled from the Virus Orthologous Groups database (vogdb.org; Supplementary Dataset 2) were detected within the proteins of the contig assemblies from the metagenomes (Supplementary Dataset 2). Phage contigs were also detected for comparison. Reads from all samples were then mapped on to these sequences for their relative abundances in each sample (Supplementary Dataset 3, see Methods), and ecological statistics held for all three metrics (MCP, TerL, contigs; Supplementary Dataset 4). Results from TerL were reported here as this was the most prevalent gene (Supplementary Dataset 4) and enabled direct comparison with prokaryotic single-genes (versus metagenome assembled genomes). Prokaryotic diversity was detected with proteins families of three genes from the Clusters of Orthologous Groups (COG) database^20^: RNA polymerase β (COG85), RNA polymerase β’ (COG86), and a ribosome-binding ATPase YchF (COG12) which has been used in a previous study^21^. Reads from all samples were then mapped on to these sequences for their relative abundances in each sample (Supplementary Dataset 3, see Methods), and ecological statistics held for all three metrics (COG85, COG86, COG12; Supplementary Dataset 4). The results of RNA polymerase β (COG85) are reported here as this was the most prevalent gene in the dataset (Supplementary Dataset 3). Details can be found in the Methods to use this approach for other datasets and studies.

**Figure 1.**
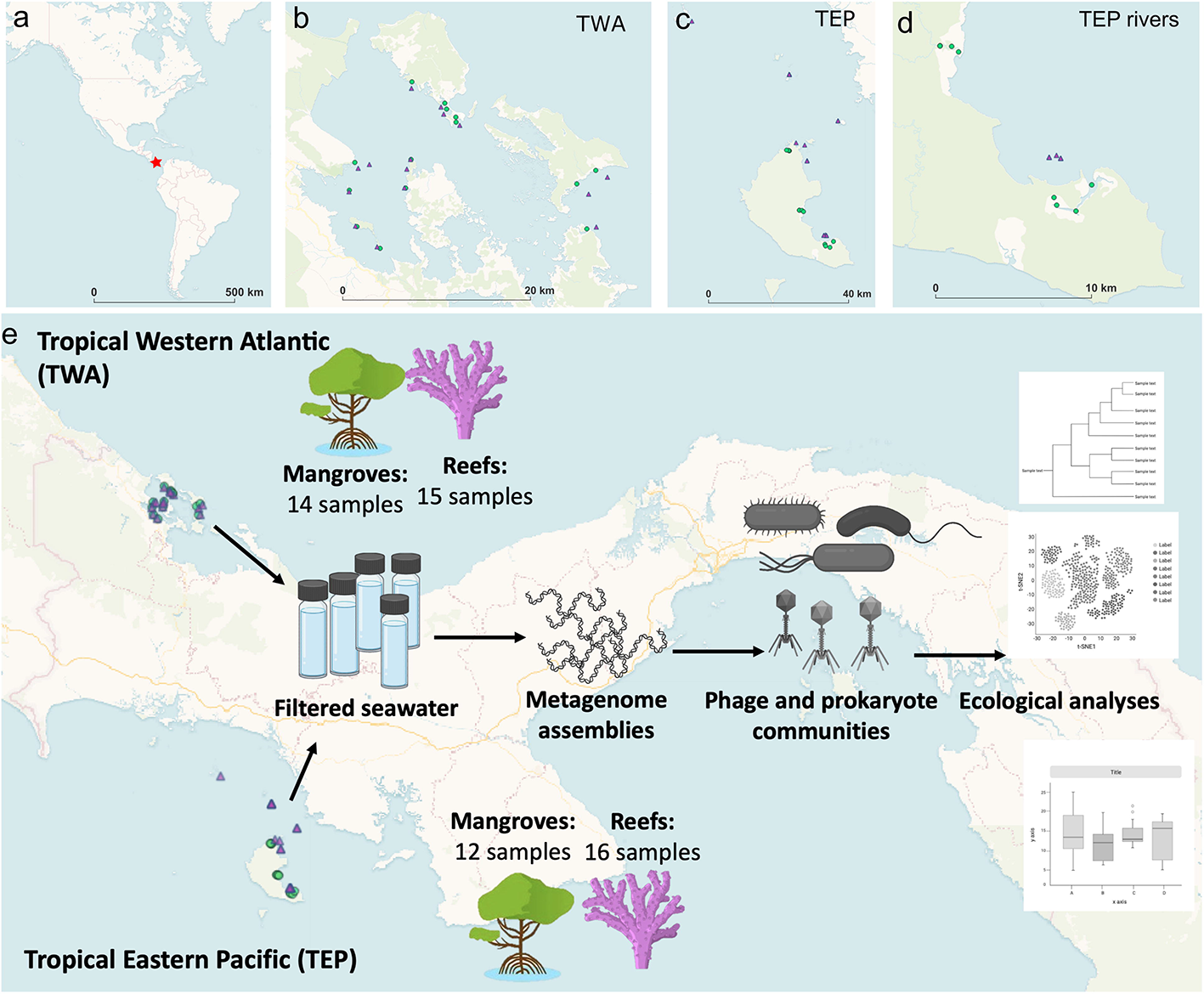
Overview of project design with maps of sample locations. (a) World map with Panama denoted as red star. (b) Map of sample sites from the Tropical Western Atlantic (TWA) coast of Panama. (c) Map of sample sites from Tropical Eastern Pacific (TEP) coast of Panama. (f) Map of TEP mangrove samples zoomed in on those collected along two freshwater rivers and the nearby reef samples. (e) Graphical abstract of project approaches. Green triangles are mangrove samples. Purple circles are reef samples.

### Proximity and physicochemical variation determine whether factors shaping phage and prokaryotic community composition align

When comparing the oceans, both phage and prokaryotic community composition significantly differed between the TEP and TWA (Figure 2a,b; ANOSIM *p values* < 0.05), and their variation correlated with each other (Mantel test *p value* < 0.05). The phage composition, however, differed to a larger extent than did the prokaryotic composition between the oceans, with only 12% of phages found in both oceans (Figure 2g) compared to 24% of prokaryotes detected in both oceans (Figure 2h). Furthermore, more physicochemical parameters varied strongly with phage composition than with prokaryotic composition (Figure 2a,b). The stronger distinction of phage composition between oceans compared to prokaryotes may result from higher dispersal limitations of most phages in the ocean. Although some phages have been detected globally^22^, this cosmopolitan distribution may be less common for phages than for prokaryotes which could be investigated further in future studies.

**Figure 2.**
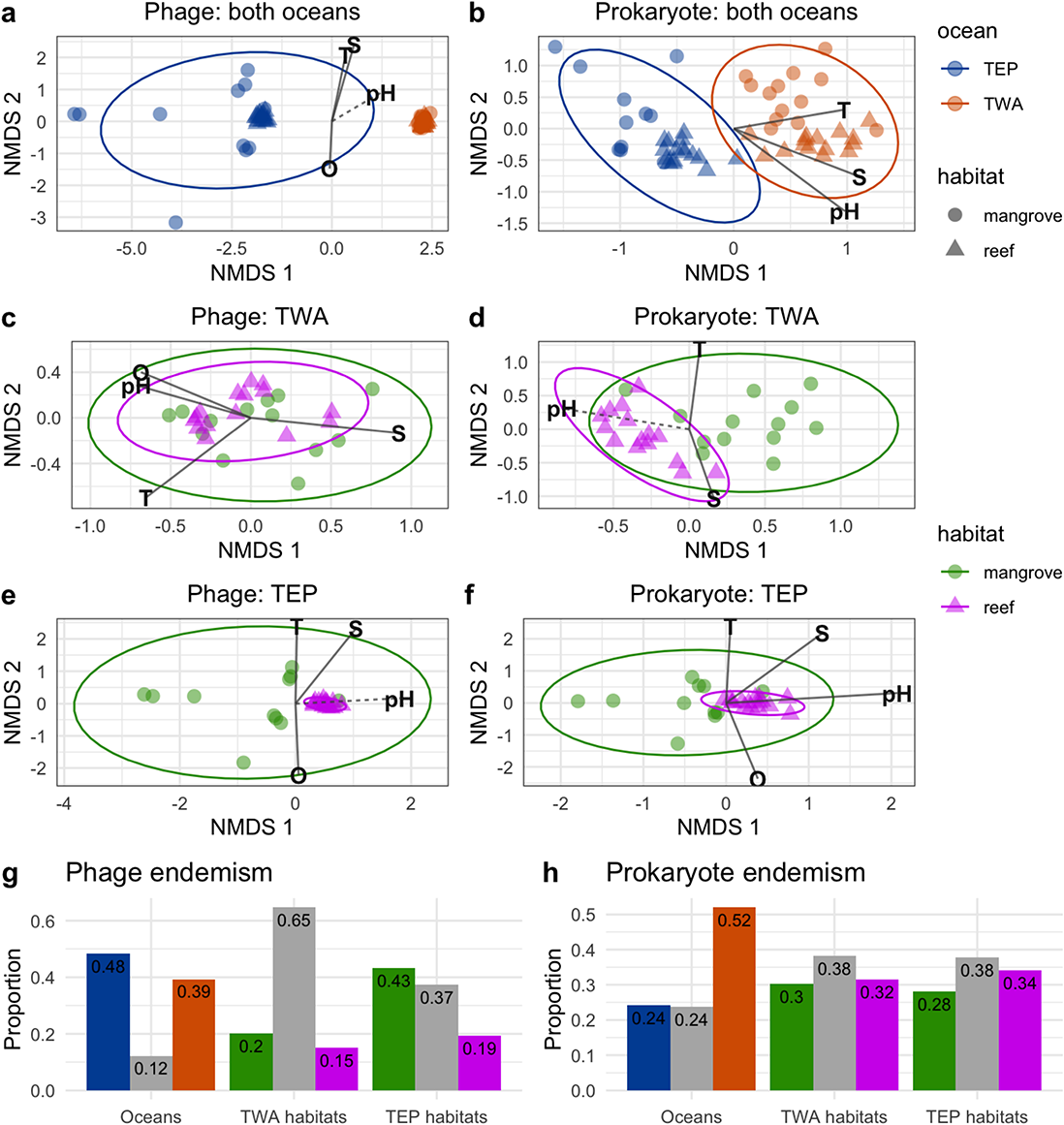
Comparisons of phage and prokaryotic community composition and endemism in marine habitats of the Tropical Western Atlantic (TWA) and Tropical Eastern Pacific (TEP). NMDS plots of samples based on phage or prokaryote community composition (Bray-Curtis distance), overlaid with environmental parameters significantly varying with community variation. Solid lines correspond to p-values below 0.01 and dashed below 0.05. Bottom barcharts compare the proportion of phages endemic to an environment or shared between them (in gray). (O - dissolved oxygen, S - salinity, T - temperature). Salinity represents total dissolved solids and specific conductivity, as these variables directly correlated with each other in this dataset.

Distinctions in community composition between the mangroves and reefs depended on the oceans. In the TEP, both phage and prokaryotic community compositions significantly differed between the habitat types (Figure 2e,f). In the TWA, only the prokaryotic composition was distinct (Figure 2d), and the phage composition did not differ (Figure 2c). In the TEP, eight of the twelve mangrove samples were collected along two rivers, with samples spanning from fully saline to fully freshwater. The other four mangrove samples were collected along a fully saline mangrove channel (Supplementary Dataset 1). Although salinity is known as one the greatest factor limiting species ranges^23,24^, the separation of the prokaryotic and phage communities here appear to cluster by river rather than by salinity (Supplementary Figure 1). Nevertheless, the same physicochemical parameters seemed to vary with the both phage and prokaryotic community composition in the TEP (Figure 2e,f). This suggests that the phage and prokaryotic communities in the TEP are likely impacted by dispersal and environmental parameters similarly. Meanwhile in the TWA, physicochemical differences between mangroves and reefs were less pronounced than in the TEP (Supplementary Figure 2), and these habitats were closer in proximity (Figure 1). Despite the lower variation in physicochemical parameters within and between reef and mangrove habitats, prokaryotic community composition partitions between habitats. This suggests that prokaryotic communities may respond to factors that we have not measured in this study, such as the distribution of dissolved and particulate organic matter. The close proximity of the mangroves and reefs, however, may have resulted in high dispersal of phages between the habitat types leading to lower distinctions in the composition, as most phages were found in both habitat types (65%; Figure 2g). The dispersal of phages between habitat types can result in a lag or delay in the shifts of phage community structure to changes in host composition because phages can only replicate upon attaching to and infecting their hosts.

Taken together, these results suggest that phage and prokaryotic community composition align when environmental conditions and spatial scales strongly structure putative host communities such as for the TEP samples (Figure 2a,e,f). Meanwhile, when these parameters are less variable, as in the TWA here, dispersal forces may structure phage communities more so than for putative host communities (Figure 2b,c,d). Another explanation between uncoupled patterns of composition in the TWA could be that physicochemical parameters interact differently on the phage and prokaryotes. For instance, pH can impact the adsorption of phages to their hosts, despite the presence of their hosts^25^. These parameters, however, would need to be tested directly.

### The most prevalent and influential phages and prokaryotes distinguishing the communities belong to diverse taxa and ecological groups

To determine which groups of phages and prokaryotes were driving the distinctions in the composition of communities, we classified the sequences using multiple approaches. The phages were classified based on the taxonomy of their putative host estimated by the alignment of the terminase large subunit (TerL) sequences to genes of RefSeq 207 and examining the host of the hits. RNA polymerase beta subunit (RNAP β) sequences used to represent prokaryotic diversity here were classified based on the consensus classification of the contig on which the RNAP β was present (See Methods for details).

Of the top ten most prevalent genera based on average relative abundance across samples, only three genera overlapped for prokaryotes and putative phage hosts: *Synechococcus*, *Prochlorococcus*, and *Pelagibacter* (Figure 3). These genera are known as dominant members of the ocean^26,27^; furthermore, because the phage sequences may also correspond to integrated phages of the prokaryotic community, this may have resulted in the co-prevalence of these genera in both phage and prokaryotic communities. Nevertheless, the general lack of overlap in prevalent phage and prokaryotic genera may have resulted from several factors such as technical limitations in classifying both the phages and prokaryotic sequences or that most viral lysis occurs for rare but highly productive microbes, as has been observed off the coast of British Columbia in Canada^28^, which would result in dominant viruses that infect rarer hosts.

**Figure 3.**
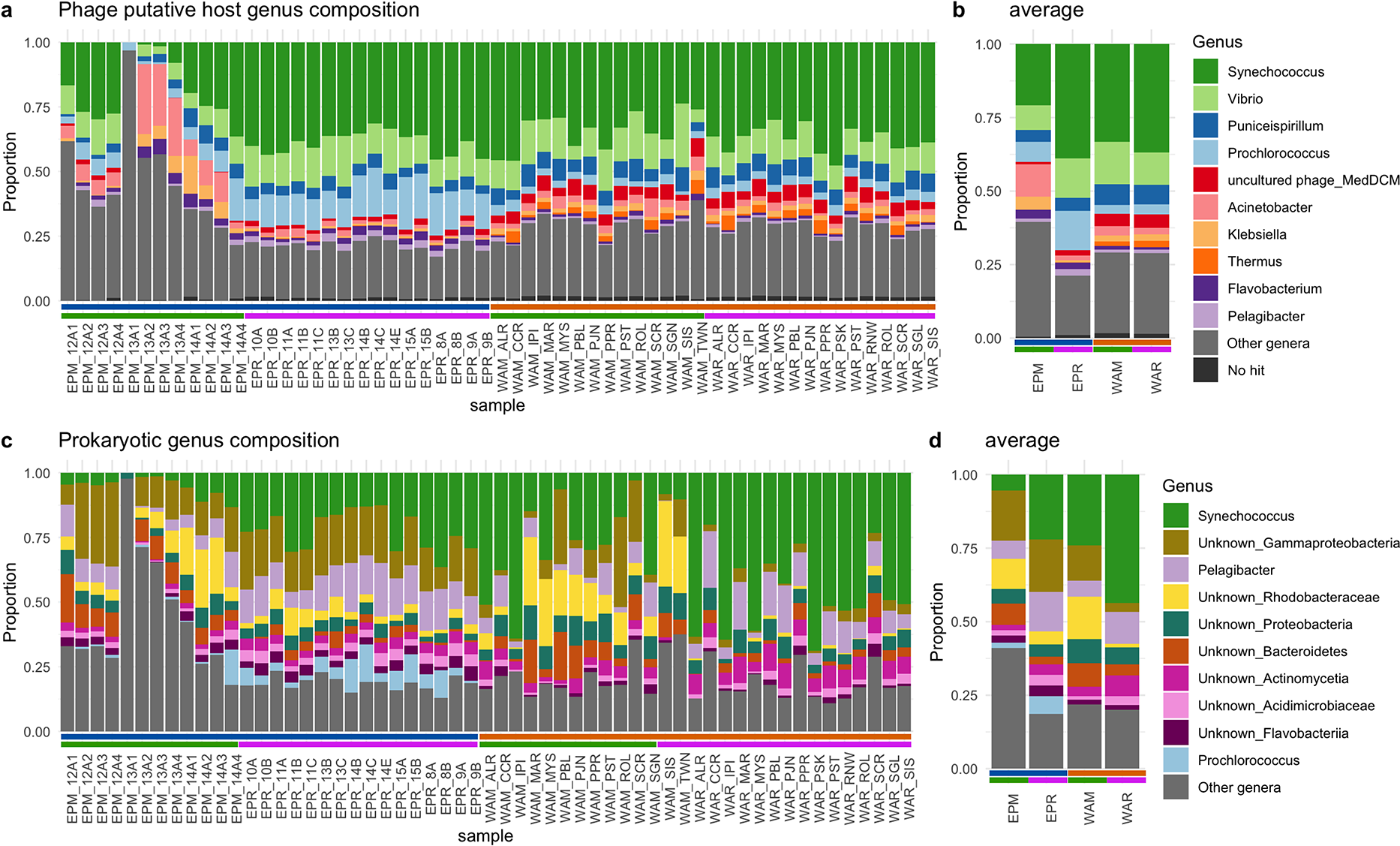
Genus composition of phage putative host communities and prokaryote communities. Stacked barplots of the phage putative host communities (a) or prokaryotic communities (c) colored and sorted by top ten average most abundant genera. Color strips on bottom indicate ocean and habitat where samples were collected: top row: navy = TEP; orange = TWA; green = mangroves; pink = reefs. (b,d) the average putative host genera composition (b) or prokaryotic genera composition (c) of ocean and habitat type combination EPM = TEP mangrove, ERP = TEP reef, WAM = TWA mangrove, WAR = TWA reef.

In general, the average genus composition of both the putative phage hosts and of the prokaryotes corroborate the compositional distinctions observed above when using sequence diversity (Figure 2), with the phage communities being highly similar between mangroves and reefs in the WA, but very distinct in the TEP (Figure 3b), and the prokaryotic communities being distinct between mangroves and reefs in both oceans (Figure 3d). In both phages and prokaryotes, the enrichment of *Prochlorococcus* in the TEP relative to the TWA highlights the physicochemical features of the ocean, as the TEP sites were more exposed to pelagic waters than the TWA sites and *Prochlorococcus* is known to be more dominant in pelagic waters than coastal waters where *Synechococcus* is prevalent^29^. Notably, a fully freshwater sample (EPM_13A, 0 ppt salinity) only contained *Prochlorococcus* of the top genera in the putative host community for the phages (Figure 4a). *Prochlorococcus* bacteria are rarely found in brackish or freshwater conditions^30,31^, and instead, a *Prochlorococcus*-like bacteria that is larger in cell size than its marine counterpart has been reported in estuaries^31^. Thus, the presence of this phage terminase with homology to that of a *Prochlorococcus* phage in the fully fresh sample suggests that either (i) this phage infects this *Prochlorococcus*-like freshwater bacteria, (ii) that it has a broad host range that enables it to infect marine and freshwater bacteria, (iii) or that its homology is a result of the limitation of the reference database. Of the prokaryotic community in this freshwater sample, only an unknown genus in the Proteobacteria phylum was found that was also prevalent in the other samples (Figure 4c), which is unsurprising as diverse Proteobacteria are common in freshwater systems^32^. The remarkable divergence of the genera in this freshwater sample for both the prokaryotes and putative host community of the phages highlights the crucial role of salinity in shaping microbial communities^23,24^.

**Figure 4.**
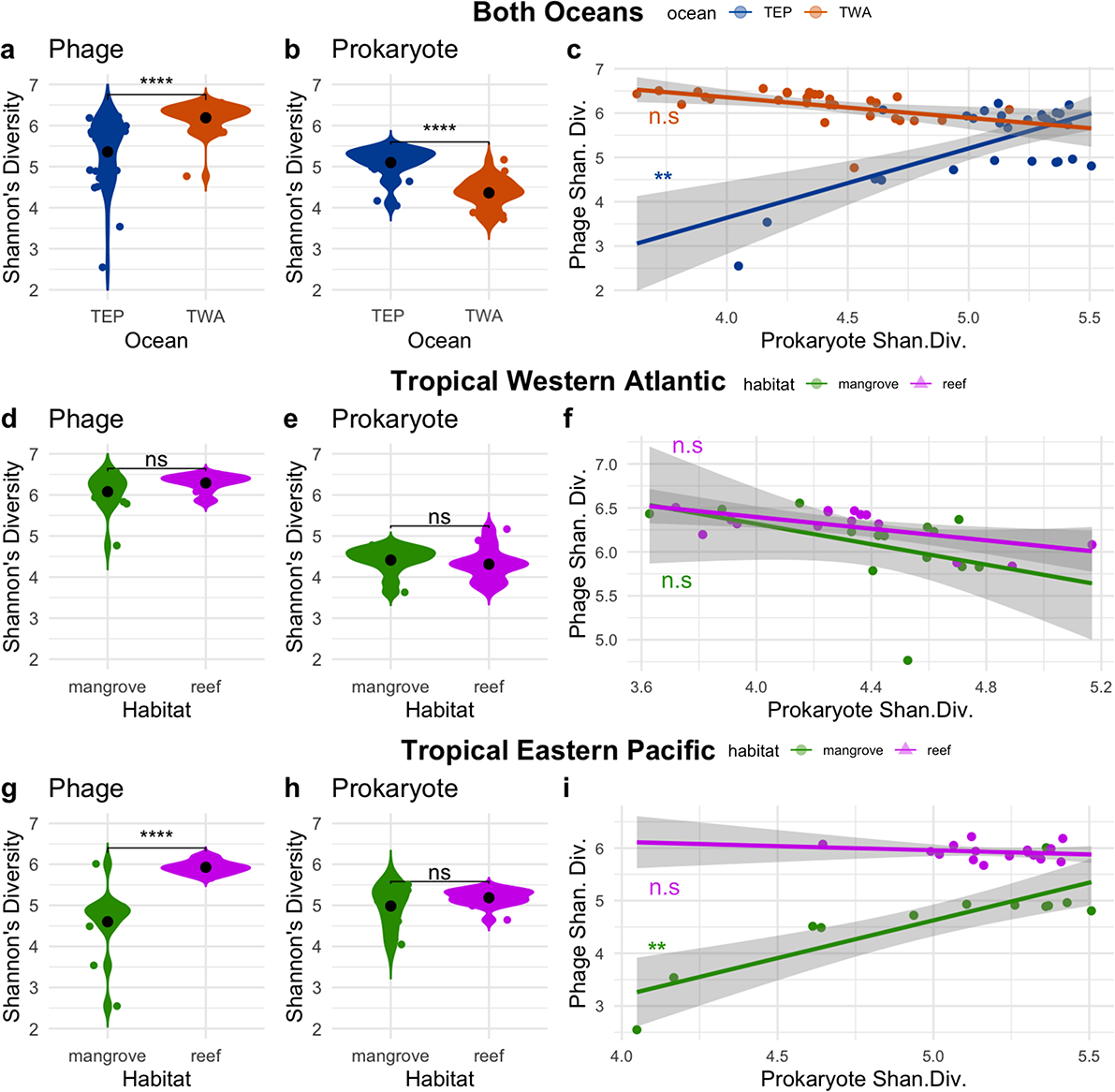
Phage and prokaryotic Shannon’s Diversity in marine habitats of the Tropical Western Atlantic and the Tropical Eastern Pacific. 2a,b,d,e,g,h are violin plots of Shannon’s Diversity of phages and prokaryotes in different samples. 2c,f,i are scatterplots of phage Shannon’s Diversity plotted against prokaryotic Shannon’s Diversity in a sample, with linear regression lines drawn and standard deviations shaded in gray. (Shan. Div. = Shannon’s Diversity).

We then examined which phages and prokaryotes drove the most variation between the samples, determined by those that significantly varied the most with variation in the communities (envfit test; *p values* < 0.05; See Methods; Supplementary Dataset 5). When examining all samples of both oceans and habitats, the phage (WA_000000419261_10), whose terminase showed high homology to that of the *Puniceispirillum phage HMO-2011*, drove the most variation followed by ten other equally influential phages that putatively infect a diversity of host genera (*Prochlorococcus, Puniceispirillum, Acinetobacter, Mycobacterium, Kiloniella, Laceyella, Escherichia*) spanning four phyla (Supplementary Dataset 5). Matching the whole community distinctions between oceans and habitats of Figure 2, all but one of these 11 phages were exclusively detected in one ocean (TEP or TWA), with phages of the TWA mostly present in both mangroves and reefs and most of those exclusive to the TEP found only in mangrove samples. In contrast, variation in the prokaryotic communities was primarily driven by *Synechococcus* bacteria (top 3 most influential; Supplementary Dataset 5), which follows its known distinction between pelagic and coastal conditions^29^, such as the TEP between TWA here. The elevated importance of phages predicted to infect chemoheterotrophic in driving phage community composition compared to the elevated importance of photoautotrophic bacteria in driving prokaryotic community composition further highlights that phage lysis predominantly occurs on the most productive members of the community, which are often heterotrophic bacteria that experience boom-and-bust cycles as nutrients become available^28^.

When examining samples of the TWA and TEP separately, the primary genera or putative host genera driving the variation in the prokaryotic and phage communities respectively aligned in trophic niche for the TWA but contrasted in trophic niche for the TEP. This is surprising because the phage and prokaryotic communities did not align in habitat distinction or physicochemical parameters driving their composition in the TWA (Figure 2c,d) but they did in the TEP (Figure 2e,f). In the TWA, the two phages that drove most of the variation showed high homology to the terminase of *Pelagibacter* phage HTVC008M and the *Puniceispirillum* phage HMO-2011, host genera that are both heterotrophic bacteria found throughout the global ocean^26,33^. For the prokaryotes, the top genera also belonged to heterotrophic groups with the top prokaryote belonging to an uncultivated genus WTJO01 in the Puniceispirillales order, and the next most influential belonging to an uncultivated genus UBA974 in the Flavobacteriales order. These heterotrophic bacteria are also found throughout the oceans^26,34^. The overlap in trophic niche of these genera for driving the phage and prokaryotic communities in the TWA, despite the differences in physicochemical and habitat distinctions, highlights the robust conditions that these groups can inhabit which could explain the lack of alignment in the environmental features driving the overall phage and prokaryotic community compositions.

In contrast to the TWA, the putative hosts of phages driving the variation of phage communities within the TEP did not align trophically despite their overlap in significant physicochemical parameters and habitat distinctions (Figure 2e,f). The most influential phages primarily putatively infect bacteria belonging to the photosynthetic *Synechococcus* genus (seven of the top ten), while the most influential prokaryotes primarily belonged to unknown genera in the Betaproteobacteria class (Supplementary Dataset 5). Although these genera contrast each other in trophic lifestyles, these bacteria are known to be highly influenced by salinity^31,35,36^, which widely varied in the TEP as the mangrove samples were collected along freshwater rivers. These results suggest that while phage and prokaryotic communities both vary substantially with salinity, the types of bacteria and putative hosts of phages that are most affected by salinity in these sites do not necessarily align.

### High prokaryotic diversity is rarely coupled with high phage diversity

Through selective killing by phages and resistance mechanisms by prokaryotes, phages and prokaryotes are known to drive each other’s evolution and microdiversity^37,38^, but how these interactions impact macrodiversity remains poorly studied. Here, we examined the alpha diversity of samples to uncover which environments contain high phage and prokaryotic sequence diversity. We used the Shannon’s Diversity index to measure alpha diversity, as this metric accounts for both richness and evenness^39^. Taxa are proxied here as unique marker sequences (see Methods). When comparing diversity between oceans, surprisingly, phage communities were significantly more diverse in the TWA (Figure 4a) while prokaryotes were more diverse in the TEP (Figure 4b). These patterns between the oceans held when comparing mangrove and reef samples separately (Supplementary Figure 3). The contrasting diversity patterns of phage and prokaryotes between oceans may be a result of several abiotic and biotic factors. Although the TWA is generally more oligotrophic than the TEP, the bay where the samples were collected in this study has historically been subject to high levels of runoff, which has been found to elevate bacterial production and density but result in decreased bacterial diversity compared to nearby pristine sites^40^. Thus, the reduced prokaryotic diversity in the TWA compared to the TEP may be due to the pollutants, while the elevated phage diversity in the TWA compared to the TEP may be due to increased bacterial production and thus phage replication and release. Alternatively, the higher phage diversity in an environment that has lower prokaryotic diversity could be because a variety of phages infect the same hosts. This would mean that a low diversity of phages could infect a high diversity of prokaryotes. The ecological conditions that would enable the coexistence of diverse phages that infect the same host may be related to the contact rates with hosts^41^ or flux between environments introducing novel phages, which may differ between the TEP and TWA, but these would need to be tested directly.

We then examined diversity within each ocean between habitat types. Within the TWA, phage communities were equally diverse between the mangroves and reefs (Figure 4d). Similarly, prokaryotic communities were equally diverse between mangroves and reefs in TWA (Fig 4e), despite significant differences in their composition between these habitat types (Figure 2d). In the TEP, phage diversity was lower in the mangroves than the reefs (Figure 4g), but prokaryotic diversity did not significantly differ between the habitat types, despite having significantly different composition (Fig 2f). This lack of alignment between the habitat types for prokaryotes in compositional variation and Shannon’s demonstrates that high diversity within samples (alpha diversity) does not always correspond to high diversity between samples (beta diversity)^42^. For example, a study by Walters and Martiny (2020)^42^ that compared the microbial diversity across a range of ecosystems found that soil samples have the highest number of microbial species (alpha diversity), but sediment, biofilms, and inland waters had the greatest variation in communities between samples (beta diversity). The conflicting alpha and beta diversity patterns of the prokaryotic communities here thus suggest that perhaps niche space is similar across habitat types leading to similar alpha diversity but competition, or local adaptation, within each habitat is strong enough to lead to distinct members of each community or strong beta diversity. Regarding the phage diversity patterns, the lower diversity of the phage communities in the mangroves than reefs in the TEP is likely due to salinity differences between these habitat types, with a median 28.93 ppt in the mangroves versus. 30.655 ppt in the reefs. Furthermore, the mangroves were more acidic than the reefs (median pH 7.885 vs 8.09). While these physicochemical differences between TEP mangroves and reef did not manifest in prokaryotic diversity differences, the phage diversity may have been impacted, as pH and salinity are known to impact adsorption rates of phages to their hosts^25,43^.

When plotting phage diversity against prokaryotic diversity, we found that phage diversity rarely correlates with prokaryotic diversity, despite the inherent link of phages to their hosts for replication. In fact, in the TWA samples, their diversities appear to negatively correlate when examining all TWA samples together and separately by habitat type, but this was not significant (Fig 2c,f). In the TEP, there is a significant positive correlation when examining the samples together (Fig 2c), but this is likely driven by the mangrove samples (Fig 2i), as there was no significant correlation between phage and prokaryotic diversity the TEP reef samples (Figure 2i). The positive correlation of phage and prokaryotic diversity in the TEP is likely due to salinity differences in the TEP mangrove samples, with two considered freshwater (~0 ppt). Upon removing these two fresh samples, the correlation of diversities is no longer significant (Supplementary Figure 4), highlighting the impact of extreme salinity differences on phage-host interactions^43^.

#### Differences in correlations with physicochemical parameters between phage and prokaryotes help explains their decoupled relationship

Because phage and prokaryotic diversity rarely correlated with each other here, we plotted phage and prokaryotic diversity against the measured physicochemical parameters of each environment to uncover other potential drivers of their diversity separately (Figure 5). Following our hypothesis that phage and prokaryotic diversity patterns will align most when environmental gradients are high, their diversities generally correlated in the same direction with most of the parameters in the TEP, which had great variation in the parameters compared to the TWA samples. For example, salinity varied widely between the mangrove samples of the TEP (0-30 ppt), and phage and prokaryotic diversity both significantly positively correlated with salinity.

**Figure 5.**
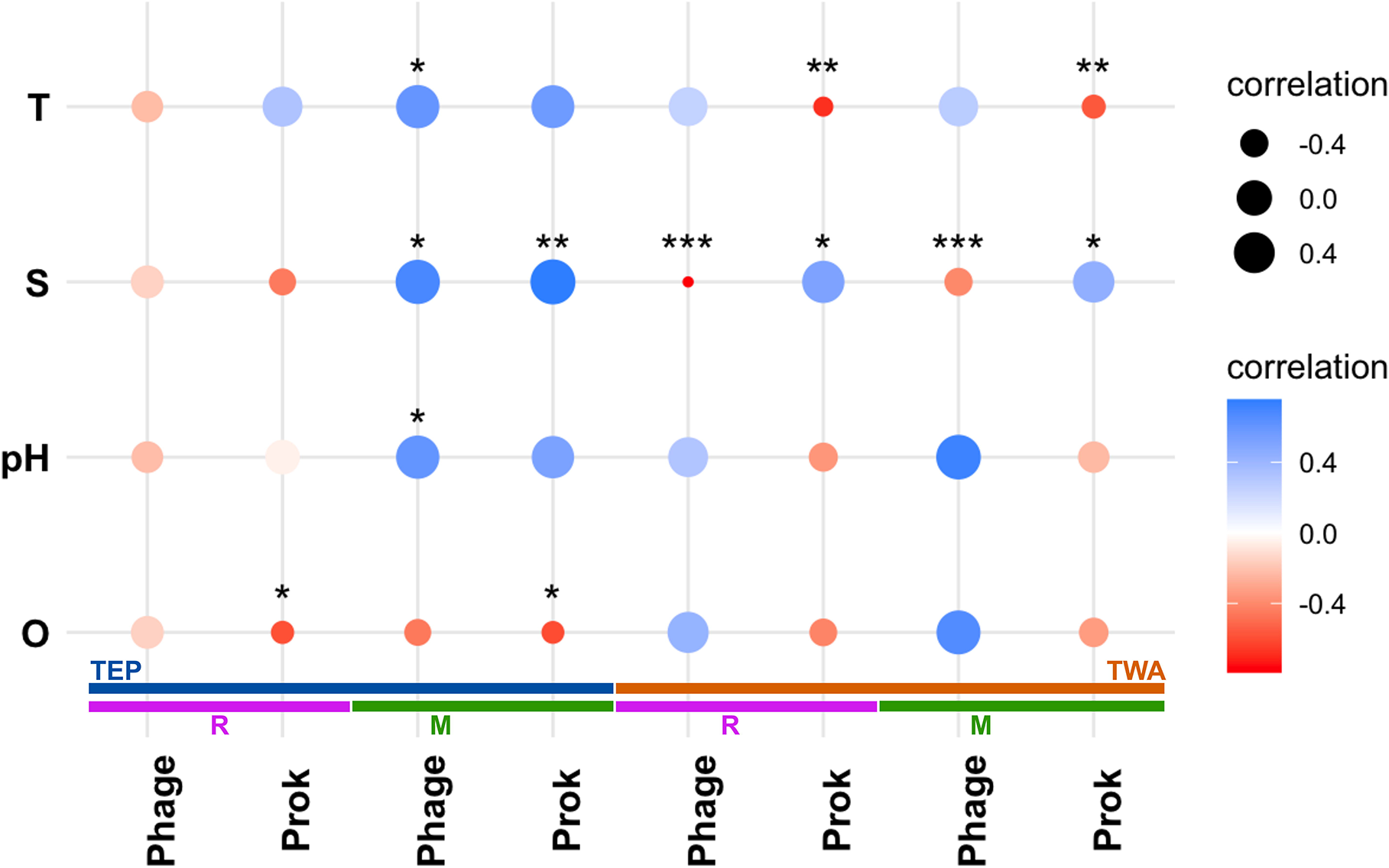
Bubble plot of correlations between measured physicochemical parameters (x-axis) and phage and prokaryotic diversity (y-axis) of the habitats in each ocean (color strips). Color and size of dot correspond to correlation strength. Stars correspond to p value significance (* < 0.05, ** < 0.01, *** < 0.001). Abbreviations: T - temperature, S - salinity, O - dissolved oxygen, Prok - prokaryote, R - reef, M - mangrove. Color strips: blue = TEP, orange = TWA, purple = reef, green = mangrove.

Meanwhile, in the TWA salinity only ranged between 32 and 34 ppt, and phage and prokaryotic diversities tended to correlate with the parameters in the opposite directions of each other. In both mangroves and reefs of the TWA, phage diversity correlated negatively with salinity while prokaryotic diversity correlated positively with salinity.

Taken together, the lack of strong relationship between phage and prokaryotic diversity (Figure 4), in addition to their inconsistent correlations with the physicochemical parameters measured here (Figure 5), exemplify the nuances in the relationship between the viral and host diversity. For instance, variation in phage host ranges or variation in host resistance to phage infections could weaken the correlation of their diversities. Furthermore, environmental variation may further decouple their diversity relationship. Our results suggest that when this environmental variation or spatial distances are relatively small between sites, such as between habitat types of the TWA, prokaryotic diversity may be impacted more than phage diversity by these local conditions. Future work could include measuring host production, phage production, and phage host ranges in isolation against different physicochemical parameters to test these hypotheses more directly.

## CONCLUSION

In this study, we leveraged the unique biogeography of the Isthmus of Panama to compare drivers of phage and prokaryotic diversity at both global scales between oceans and local scales between habitat types within each ocean by examining mangrove and reef habitats of the TEP and TWA coasts (Figure 1). We found that drivers of phage and prokaryotic communities align most when physicochemical and spatial scales are sharp, such as between the oceans and between the TEP mangroves and reefs. Meanwhile, these factors diverge when there are subtle physicochemical differences and minimal physical separation in environments, like between the mangroves and reefs of the TWA. In these cases, prokaryotic communities may locally adapt to the minor environmental differences, as we observed distinction between prokaryotic communities of the mangroves and reefs. The phage communities, however, may be influenced more by high dispersal between the environments, overwhelming environmental or habitat distinctions, as we observed no significant difference between mangroves and reefs of the TWA. A similar pattern has been observed in a freshwater spring system of southern Florida, where the prokaryotic communities were distinct between the river, head, and mixed zones, but the phages communities were not distinct between the head and mixed zone, which the authors attributed potentially to high dispersal of phages between these two zones^14^.

Despite cases when drivers of phage and prokaryotic community composition align, our results show that putative host genera of phages that drive phage communities differ from prokaryotes in all spatial and physicochemical scales. Very few of the most dominant phage members infect genera of the most dominant prokaryotes. This may be because most phages are infecting the most productive prokaryotes which exhibit boom and bust reproductive cycles, rather than the most stably abundant^28^. This infection pattern would support the popular phage-host interaction model, the Kill-the-Winner model, in which phages rise in abundance to kill the most dominant prokaryotes^44^. However, deviations from the Kill-the-Winner model have been observed in a freshwater lake where the abundance of some phages have been found to peak before, during, or after their host’s peak in abundance^45^, and we would thus need time series data to resolve this possibility.

Counterintuitively, we found that high phage diversity is rarely coupled with high host diversity. The only context when their diversity correlated was in the TEP mangrove samples when fully freshwater samples were included, which points to the strong role of salinity in shaping both prokaryotic and phage communities^23,43^. Because phage diversity was higher in the TWA than the TEP, we suspect that host production rates may drive phage diversity such that even if there are blooms of a single bacteria, a variety of phages may surface to infect that host. Conversely, a lower phage diversity amidst high prokaryotic diversity may result if phages have broad host ranges, which may be related to host contact rates^41^. We also saw that phage and prokaryotic diversity can be driven by physicochemical parameters differently and align the least when environmental variation and physical separation is subtle, such as in the TWA sites, compared to when they are more pronounced, such as in the TEP sites. Together, these trends indicate that the strength of the link of phage communities between potential host communities depends on the level of variation in the environment. Future work that includes additional measurements such as phage production, prokaryotic growth, organic matter concentrations, and more could reveal more precise dials in the constraints on the link in patterns of phage communities to those of prokaryotic communities.

All in all, this study provides a framework and demonstrates an application for comparing phage and prokaryotic community composition and diversity in a variety of marine environments. We uncover conditions when the tight links of phages and prokaryotes result in similar factors driving their diversity and composition, such as between oceans, and when these tight links are weakened, such as between adjacent but distinct habitat types. By understanding when these phage-host links are strengthened or weakened, we can better predict the outcome of interactions between phages and prokaryote populations of different environments to inform models of nutrient cycling mediated by microbes and the release of organic matter through viral lysis of microbes.

## METHODS

### Sample and environmental data collection

Seawater samples were collected ~1m above the seafloor on coral reefs and mangroves (1-4m depth) in the TEP and TWA coasts of Panama in 2017 (see Supplementary Dataset 1 for coordinates and collection dates). Seawater samples were collected in sterile Whirl-Pak Bags and kept on ice and in the dark until filtration at either the Smithsonian Tropical Research Institute (STRI) Coiba (TEP) or Bocas del Toro research stations (TWA), where they were then vacuum filtered through 0.22 μm nitrocellulose membranes (Millipore). Filters were frozen and transported to STRI’s molecular facility at Isla Naos Laboratory in Panama City in liquid nitrogen and stored at −80 °C until DNA extractions. DNA was extracted from each filter using a Qiagen Powersoil extraction kit following the manufacturer’s protocol with minor modifications to increase the yield46. Metagenomic shotgun libraries were prepared with the Illumina DNA Nextera Flex kit following the manufacturer’s protocol. Shotgun metagenomics reads were sequenced on an Illumina Nextseq platform. Dissolved oxygen, temperature, salinity, and pH were measured with a pre-calibrated Professional Plus handheld YSI (Yellow Springs, USA).

### Metagenome preparation, sequencing, and assembly

We used Trimmomatic (v0.39)^47^ for adapter clipping and initial quality trimming of raw metagenomic data (N = 57). We used anvi’o (v7.1)^48^ to build a Snakemake (v.5.10.0)^49^ workflow for co-assembly analysis. In the workflow, we used iu_filter_quality_minoche from the Illumina Utils package (v2.12)^50^ for additional quality filtering and MEGAHIT (v1.2.9)^51^ for co-assembly (– min-contig-len: 1000, –presets: meta-sensitive). We performed three separate co-assemblies using MEGAHIT based on initial assessment of the metagenomic data. All TWA samples (reef and mangrove) were co-assembled (n = 29); from the TEP, we performed one co-assembly for reef samples (n = 16) and another for mangrove samples (n = 12). Next, we used anvi-gen-contigs-database to generate a database of contigs. Within the Snakemake workflow, KrakenUniq (v0.5.8)^52^ was used for taxonomic classification of short reads against a user-constructed database of archaea, bacteria, viral, fungi, and protozoa reads from RefSeq and the NCBI nt database. Taxonomic classification of contigs was performed using Centrifuge (v1.0.4_beta)^53^, against the bacterial, archaeal, human, and viral genomes database.

### Phage marker gene and contig curation

For the marker gene detection, open reading frames (ORFs) were predicted with prodigal^54^ (-p meta -a -d) on contigs of all sizes (753,612 EP; 574,304 WA contigs | 2,168,906 EP; 1,756,476 WA ORFs; 3,925,382 total ORFs). Amino acid sequences of the ORFs were then searched against all MCP and TerL HMM profiles available in Virus Orthologous Group database (vogdb.org) version 208 (Supplementary Dataset 2) using hmmsearch (hmmer.org; E value < 0.00001, bitscores > 41 and > 33, respectively, minimum length of open reading frame >=826 and >= 885 nucleotides, respectively). The threshold bitscores were determined by searching proteins predicted with prodigal (default per genome) from all *Caudovirales* genomes from Viral Genomes Portal downloaded on July 26, 2021 against the MCP and TerL profiles, taking the top hit from each genome and identifying the minimum bitscore required to include at least 98% of hits. After filtering for bitscore, the minimum length of a hit was decided based on containing at least 98% of those reference hits. This resulted in 3,749 MCP genes and 5,369 TerL genes.

These were then de-replicated at 100% identity across the entire length of one sequence using BLASTn^55^, which resulted in 3,722 representative MCP and 5,350 TerL (See Data Availability). For the detection of phage contigs, contigs over 10 kilobases (7,619 EP; 10,839 WA) were run through VirSorter2^56^ and CheckV^57^ as follows. First, contigs over 10 kilobases (EP: 7,619, WA: 10,839;) were run through VirSorter2 (virsorter run --min-score 0.5 all) and retained if they scored over 0.5 for dsDNAphage as their max_group (EP: 1,513, WA: 3,272). These contigs were then run through CheckV (checkv end_to_end) to trim potential host genomes flanking the contigs. Trimmed provirus and virus sequences were combined and filtered for at least 10kb (EP: 1,482, WA: 3,203). The trimmed sequences were then run through VirSorter again and retained if they scored over 0.95 or scored at least 0.5 and encoded at least 2 phage hallmark genes. This resulted in 3,885 contigs. Virus detection summary for each contig is in Supplementary Dataset 2.

### Prokaryote marker gene curation

The same ORF and amino acid sequences used for the phage marker gene detection were searched against HMM profiles corresponding to genes to the Clusters of Orthologous Groups (COG) protein families of COG0012 (COG12, ribosome-binding ATP-ase), COG0085 (COG85, RNA polymerase β subunit), and COG0086 (COG86, RNA polymerase β’ subunit)^20^ jointly using hmmsearch (E value < 0.00001, bitscores cutoffs of 210, 200, and 200 respectively^58^. See Supplementary Dataset 4 for the number of hits of each gene.

### Distribution detection

Reads from all samples were subset to an even depth to the number of reads in the sample with the fewest reads (2,992,107 reads) with seqkit^59^ sample (−s 1000, −2). Reads were then mapped to an index of the phage marker genes, phage contigs, and prokaryote marker genes made with minimap2^60^ -x sr. CoverM^61^ (https://github.com/wwood/CoverM) was then used for the mapping (coverm contig --min-read-percent-identity 95 -m covered_fraction rpkm count variance length -- minimap2-reference-is-index --min-covered-fraction 0 --coupled) and retained with 50% gene covered or 20% of contig covered^62^ (Supplementary Dataset 3). See Supplementary Dataset 4 for the number of each sequence type detected in at least one sample.

### Visualizations, statistical analyses and sequence benchmarking

All plots aside from the maps were created in R (version 3.5.1)^63^ with Rstudio (version 1.1.456)^64^ using vegan^65^, ggpubr^66^, and ggplot2 (3.1.1)^67^. Maps were created with QGIS (3.24) using the Voyager plug-in for the base and overlaid with sample data. Because statistics and trends held regardless of protein examined per bacteria or phage (Supplementary Dataset 4), we focused on the TerL results to represent phage diversity and COG85 results to represent bacterial diversity, as these genes were the most prevalent in the dataset. Influential sequences and physicochemical parameters were identified by those varying the most with variation in the communities of all samples based on significant vector length (vegan package function envfit, perm=999, na.rm=TRUE; calculated with |NMDS1-NMDS2|; *p values* < 0.01). Community composition of samples were compared and visualized in non-metric dimensional scaling (NMDS) plots using Bray-Curtis distances of relative abundances calculated with reads per kilobase per million (RPKM) using vegan (metaMDS(distance = “bray”)). Two outlier samples were excluded in the community compositional analyses as these were highly divergent (WAM_TWN and EPM_13A1) and skewed the results (Supplementary Figure 4, 5). WAM_TWN was sampled in a highly polluted site, and EPM_13A1 was sampled from a completely freshwater sample, which likely resulted in their aberrant community compositions at the genus-level (Figure 3c,d). Significant distinctions between oceans and habitat types were determined with ANOSIM test (vegan package) based on Bray-Curtis dissimilarity matrices using the RPKM data (anosim(distance=”bray”,permutations=9999)).

### Gene taxonomy

Prokaryotic sequences corresponding to COG85 were classified via Centrifuge^53^. For the phages, amino acid sequences of TerL genes were aligned to RefSeq 207 with LAST^68^ (lastal -m 10 -f BlastTab; E value cutoff 10^−5^), and the taxonomy of the hit’s host was reported (i.e. a hit to a *Prochlorococcus* phage meant the taxonomy of *Prochlorococcus* was reported). The top hit was detected based on percent identity. The top 10 genera based on average relative abundance across samples was reported.

## DATA AVAILABILITY

Reads from metagenomes will be deposited on the European Nucleotide Archive upon publication. Sequences of marker genes and phage contigs can be found on the FigShare repository upon publication, along with the VOG and COG HMM profiles used for marker gene detection.

## CODE AVAILABILITY

Custom scripts used for this study are found in the GitHub repository (https://github.com/scubalaina/panama_phages).

## SUPPLEMENTARY FIGURES

Supplementary Figures can be found at the link here: https://github.com/scubalaina/panama_phages/blob/main/Supplementary_Figures.pdf.

## ACKNOWLEDGEMENTS

We thank members of the Aylward Lab for helpful feedback. We thank the Smithsonian Tropical Research Institute staff at the Bocas del Toro and Naos stations. This work was performed using compute nodes available at the Virginia Tech Advanced Research and Computing Center and on the Smithsonian High-Performance Cluster (SI/HPC), Smithsonian Institution (doi:10.25572/SIHPC). This work was supported by grants from the Gordon and Betty Moore Foundation awarded to STRI and UC Davis (doi:10.37807/GBMF5603), the NSF CAREER award (IIBR-2141862) to FOA and a Simons Early Career Award in Marine Microbial Ecology and Evolution to FOA. ARW was supported by an ICTAS Doctoral Scholars Fellowship. Research permits were provided by the Autoridad Nacional del Ambiente de Panamá.

## AUTHOR CONTRIBUTIONS

JJS and ML collected the samples for sequencing and the associated metadata; they also processed the samples and sent them for sequencing. JJS performed initial sequence data processing, quality control, and assembly. ARW and FOA designed the data analysis. ARW performed the data analysis and developed the manuscript. All authors contributed to the interpretation and writing of the manuscript.

## Notes

Funding Support: This work was supported by grants from the Gordon and Betty Moore Foundation awarded to STRI and UC Davis (doi:10.37807/GBMF5603), the NSF CAREER award (IIBR-2141862) to FOA and a Simons Early Career Award in Marine Microbial Ecology and Evolution to FOA. ARW was supported by an ICTAS Doctoral Scholars Fellowship.

https://github.com/scubalaina/panama_phages

### Competing Interest Statement

The authors have declared no competing interest.

